# Test-Retest Reliability of Motor Evoked Potentials Across Eight Bilateral Lower-Limb Muscles

**DOI:** 10.64898/2026.08.26.747367

**Authors:** Karly Willson, Helia Mojtabavi, Jonathan R. Wolpaw, Russell L. Hardesty

## Abstract

**Objectives:** Transcranial magnetic stimulation (TMS) is widely used to probe corticospinal excitability by eliciting motor evoked potential (MEP)s in targeted muscles, with MEP characteristics such as magnitude and latency reflecting the physiological state of the pathways being stimulated. Although numerous studies have examined MEP reliability in upper extremity muscles, less is known about the reliability of this measurement across the lower extremity. We hypothesized that inter-session, test-retest reliability of MEPs recorded simultaneously from multiple lower-limb muscles, from a single TMS location, would differ by muscle, stimulation intensity, and quantification method.

**Materials and Methods:** Ten healthy participants (5 males, 5 females) completed three TMS sessions separated by one week. At each session, the stimulation hotspot was identified using a five-location virtual grid anchored at the vertex, with electromyography (EMG) recorded from all eight muscles of interest at each grid location; the grid location producing the largest and most consistent MEPs in the tibialis anterior (TA), the primary target muscle, was selected as the stimulation site and held constant across all three sessions. MEPs were then recorded bilaterally from the TA, soleus, rectus femoris, and biceps femoris muscles at two stimulation intensities (110% and 120% resting motor threshold (RMT)). MEP size was quantified using mean rectified magnitude and peak-to-peak amplitude, and inter-session reliability was assessed using intraclass correlation coefficients (ICC). Bland-Altman analysis was used to characterize the range of measurement variability across all eight muscles.

**Results:** MEP size differed across sessions, and reliability varied by muscle, intensity, and quantification method. The highest reliability was observed in the right TA, the muscle used to establish the stimulation hotspot, using mean rectified magnitude at 120% RMT. Reliability was comparatively lower in the seven non-target muscles recorded from the same fixed stimulation site, indicating that MEP consistency was not uniform across the lower-limb musculature.

**Conclusions:** MEP reliability in the lower extremity varies by muscle, stimulation intensity, and quantification method. MEPs were most reliable in the TA, the muscle used to establish the stimulation site, while reliability was lower in the other seven muscles.

## Introduction

Transcranial magnetic stimulation (TMS) is a noninvasive procedure to stimulate cortical neurons from outside the skull, evoking motor activity in extremity muscles reflected in electromyography (EMG) signals [1]. TMS of the motor cortex causes activation of pyramidal neurons that are not currently in their refractory period (i.e., excitable neurons) to reach their firing threshold and generate a volley of action potentials that propagate down corticospinal projections to a target muscle, generating a motor evoked potential (MEP) [1, 2]. MEPs elicited by TMS serve as a marker of corticospinal excitability by indexing how easily descending corticospinal projections activate lower motoneurons [3].

MEP measurements have been extensively validated for reliability when targeting single muscles. Upper extremity muscles show consistently high reliability in both young and older adults [4, 5, 6, 7]. Similarly, recordings from the tibialis anterior (TA) demonstrate robust reliability in healthy individuals, specifically when elicited from higher TMS intensities with a controlled level of background muscle activation [8, 9, 10, 11]. Reliability has also been demonstrated in recordings from proximal lower limb muscles, including the vastus medialis, vastus lateralis, and rectus femoris [12, 13, 14, 15, 16]. However, MEP measurements from clinical populations, e.g., chronic stroke or spinal cord injury, show diminished reliability in comparison to healthy controls in the TA and in quadriceps components including vastus medialis and lateralis [9, 17, 18].

A key distinction between eliciting MEPs in muscles of the lower *vs*. upper extremities lies in their anatomical representations in the motor cortex. Cortical representations of lower limb muscles reside deeper within the longitudinal fissure compared to their upper limb counterparts. This anatomical difference poses technical challenges for lower limb TMS, as the deeper location and limited focality of the magnetic field can result in simultaneous activation of multiple lower limb muscles, complicating the isolation of specific muscle responses [19, 20]. Despite evidence that TMS applied to a single cortical location can elicit MEPs across multiple lower extremity muscles, the test-retest reliability of such simultaneous multi-muscle recordings remains poorly understood. This gap is particularly relevant given the practical advantages of obtaining comprehensive lower limb assessments from a single stimulation site.

The reliability of MEP measurements from multiple muscles stimulated at a single TMS location may be influenced by inter-muscle differences in corticospinal excitability. [21] investigated this by comparing corticospinal excitability across five unilateral lower limb muscles (TA, soleus, rectus femoris, biceps femoris, and abductor hallucis). Their findings revealed significant variations in both MEP amplitude and resting motor threshold (RMT) across muscles, with the biceps femoris exhibiting substantially higher RMT compared to the rectus femoris, soleus, and TA.

Beyond anatomical factors, measurement methodology can also influence MEP reliability. Evidence regarding optimal stimulation intensity remains mixed, while some studies report greater reliability at higher intensities [22, 23], others find no intensity-dependent differences [13]. Current guidelines recommend targeting the linear portion of the stimulus-response curve, where MEP amplitude scales predictably with intensity, thereby enhancing measurement consistency [24]. MEP amplitude can be quantified as either peak-to-peak amplitude (maximum waveform excursion) or mean rectified magnitude (integrated area under the rectified waveform). While some studies reported equivalent reliability between these quantification metrics [23], others suggest that mean rectified area yielded lower variability in MEP measurements [25].

The present study examines whether methodological factors —stimulation intensity (110% vs. 120% RMT), quantification methods (peak-to-peak amplitude vs.mean rectified magnitude), and coil positioning accuracy— influence the test-retest reliability of MEP measurements across eight lower extremity muscles—bilateral TA, soleus, rectus femoris, and biceps femoris—using a single optimized TMS location.

## Methods

### Ethics Statement

All procedures were performed at the Samuel S. Stratton VA Medical center (SSSVAMC) in Albany, NY and were pproved by the Institutional Review Board consistent with the standards of the Declaration of Helsinki. All participants provided written consent to participate in the study.

### Human Participants

The current study consisted of ten healthy participants (5 males, 5 females, age: 44. 4 ± 15. 36). All participants completed the full study duration. Inclusion criteria required participants to be age 18 or older, have no history of major neurological disease or other major medical disorders, exhibit reasonable expectation that any current medication would not change over the study period, and demonstrate an understanding of study instructions and provide informed consent. Exclusion criteria for participants included pregnancy, metal implants in or above the chest (i.e. pacemaker, cochlear implant), any history of seizure, and any damage to ear anatomy.

### Experimental Design

We employed a within-subjects design with three sessions, examining eight muscles bilaterally (TA, soleus, rectus femoris, and biceps femoris) at two stimulation intensities, 110% and 120% RMT, with 25 pulses delivered per intensity. This dual-intensity design follows established TMS reliability protocols using 110% and 120% RMT to characterize corticospinal excitability [26]. The primary dependent variable was MEP size, quantified using mean rectified magnitude and peak-to-peak amp itude.

### TMS Procedures

The overall experimental procedures are outlined in Figure 1. TMS was delivered using a Magstim device with a double cone coil over the motor cortex, targeting the right TA muscle (see Stimulation Location below for the grid selection procedure). BrainSight neuronavigation software and a Polaris motion capture camera were used to scale a virtual model of each participant’s scalp in order to maintain stimulation location accuracy and consistency. Evoked Potential Operant Conditioning System software (EPOCS) [27], a custom-made software for operant conditioning of spinal reflexes, was configured to send TTL pulses at a 6-s inter-stimulus interval (ISI) to ensure neurons of descending motor pathways exited the refractory period and were capable of firing. After each ISI, EPOCS confirmed that the participant was at rest and issued a TTL pulse to trigger the Magstim device to deliver a TMS pulse. The BrainSight system recorded the position and orientation of the coil at the time of stimulation.

**Figure 1:**
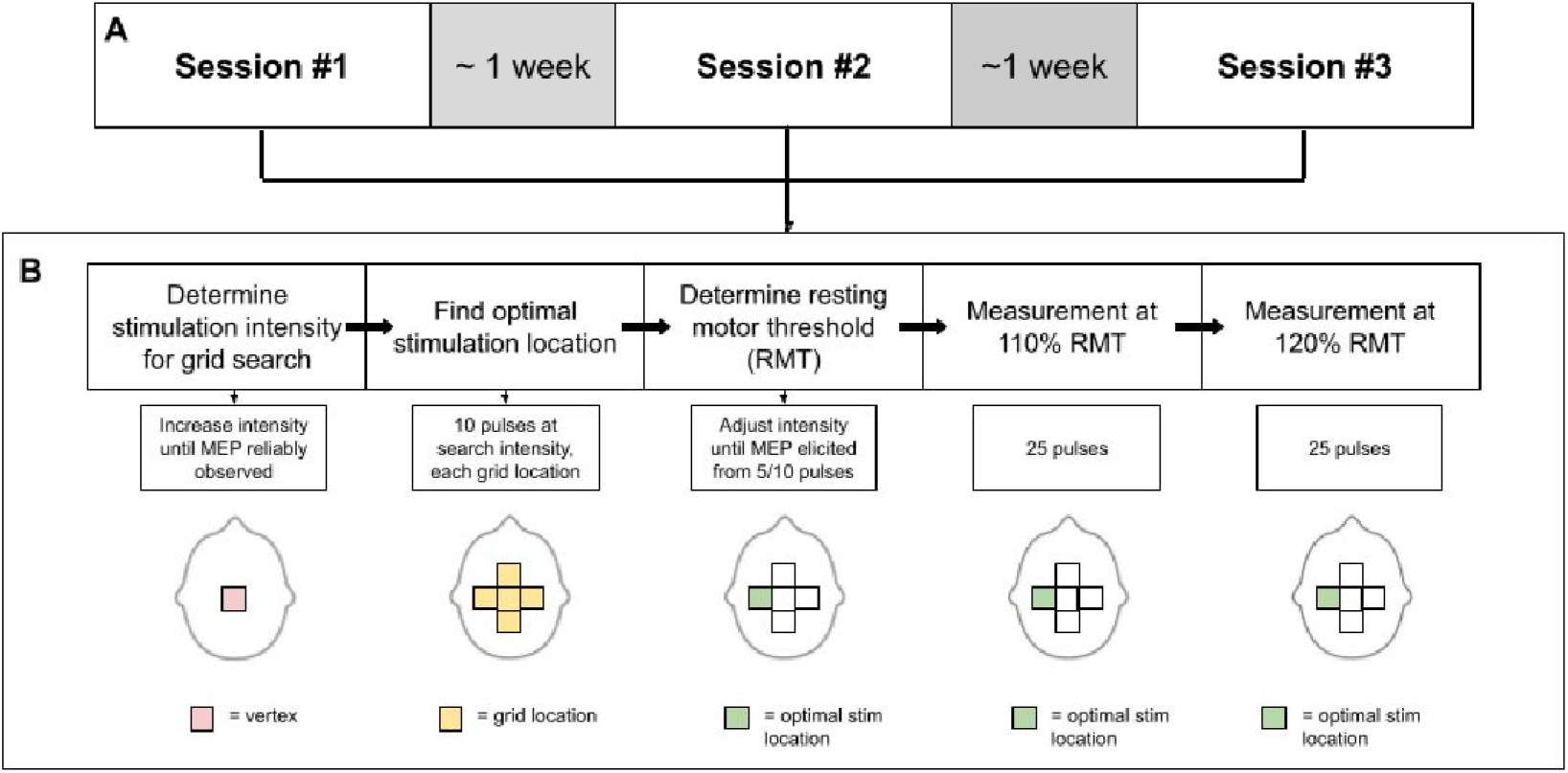
Experimental Procedures. Panel A outlines the overall structure, which consisted of three identical sessions conducted at least seven days apart. Panel B outlines the procedure followed during each of the three sessions. The grid search stimulation intensity was first determined over the vertex, represented by the pink square. At this intensity, EMG was recorded from all eight muscles of interest, with the TA prioritized as the primary target muscle, as 10 pulses were delivered at each of the five grid locations, represented by yellow squares. Once the optimal location was determined, as re resented by the green square, resting motor threshold was determined by adjusting stimulation intensity until MEPs were elicited in the target muscle from five out of ten pulses. Lastly, 50 total pulses were delivered at the optimal stimulation location (25 pulses at 110% RMT and 25 pulses at 120% RMT) while EMG was recorded from all eight muscles of interest. All diagrams not drawn to scale.

Participants returned to the laboratory at the same time of day for three successive sessions, separated by approximately one week (*M* = 9.75 days ± 5.99). At the start of each session, surface EMG electrodes were placed bilaterally over the TA, soleus, rectus femoris, and biceps femoris muscles. During the first session, each electrode location was carefully marked with skin-safe surgical markers or henna to ensure consistent EMG recording across subsequent sessions. Participants were then seated in a comfortable chair facing the Polaris motion-capture camera, which had been calibrated to the TMS coil. A headband equipped with motion-capture sensors was positioned on the participant’s head, ensuring visibility to the camera system. Using the BrainSight neuronavigation system, approximately 54 points across the participant’s skull were digitized with a sensor-equipped pointer. This digitization process allowed the system to scale a generic head model to closely match each participant’s actual head dimensions. The cranial vertex was manually determined by measuring the midpoint between the nasion and inion, and the midpoint between the right and left tragus across the skull surface. This vertex location was then registered into the BrainSight system to serve as an anatomical reference point for laying our virtual grid of stimulation locations.

#### Stimulation Location

The optimal stimulation location was defined as the area over the cortex which produced the largest and most consistent response in the target muscle [28]. The TA was defined as the primary target muscle in the present study. TMS locations were arranged in a five-location virtual grid anchored at the vertex. Two locations were positioned along the midline, 0.5 cm and 1 cm posterior to the vertex, respectively. The remaining two locations were positioned 0.5 cm to the left and 0.5 cm to the right of the point 0.5 cm posterior to the vertex, along the mediolateral axis. The BrainSight software was used to guide TMS coil placement and orientation over the scalp at these five locations. The coil was initially positioned over the vertex to deliver test stimulations at various intensities to identify an appropriate stimulation level that consistently elicited MEPs. At this stimulation intensity, 10 pulses separated by 6 s were delivered at each of the five grid locations, and EMG was recorded from all eight muscles of interest. Peak to peak MEP amplitude for each muscle were calculated and displayed per grid location using custom Python software. Two investigators reviewed this output and jointly selected the optimal stimulation location, prioritizing the grid location that produced the largest and most consistent MEPs in the TA while also accounting for responses recorded in the remaining seven muscles. Once identified, this location was held constant for the subsequent sessions.

#### Resting Motor Threshold

After identifying the optimal stimulation location, RMT was established by systematically adjusting the stimulator output until stimulation elicited MEPs on five out of ten pulses in the target muscle, i.e., the TA. RMT was defined as the lowest stimulation intensity required to produce MEPs (>50 *µ*V) in the target muscle at rest [1].

#### TMS Measurement

Participants received 50 TMS pulses over the optimal stimulation site: 25 pulses at 110% of RMT and 25 pulses at 120% of RMT, with a 6-s inter-pulse interval. Participants were instructed to remain at rest throughout all stimulation trials. Differential EMG signals were recorded continuously from all muscles of interest, amplified (gain=500), bandpass filtered (10-500 Hz), and digitized (National Instruments DAQ system). Data acquisition and storage were managed through EPOCS software. MEPs were extracted using a post-stimulus window of 10-80 ms following each detected stimulation, as illustrated in Figure 2.

**Figure 2:**
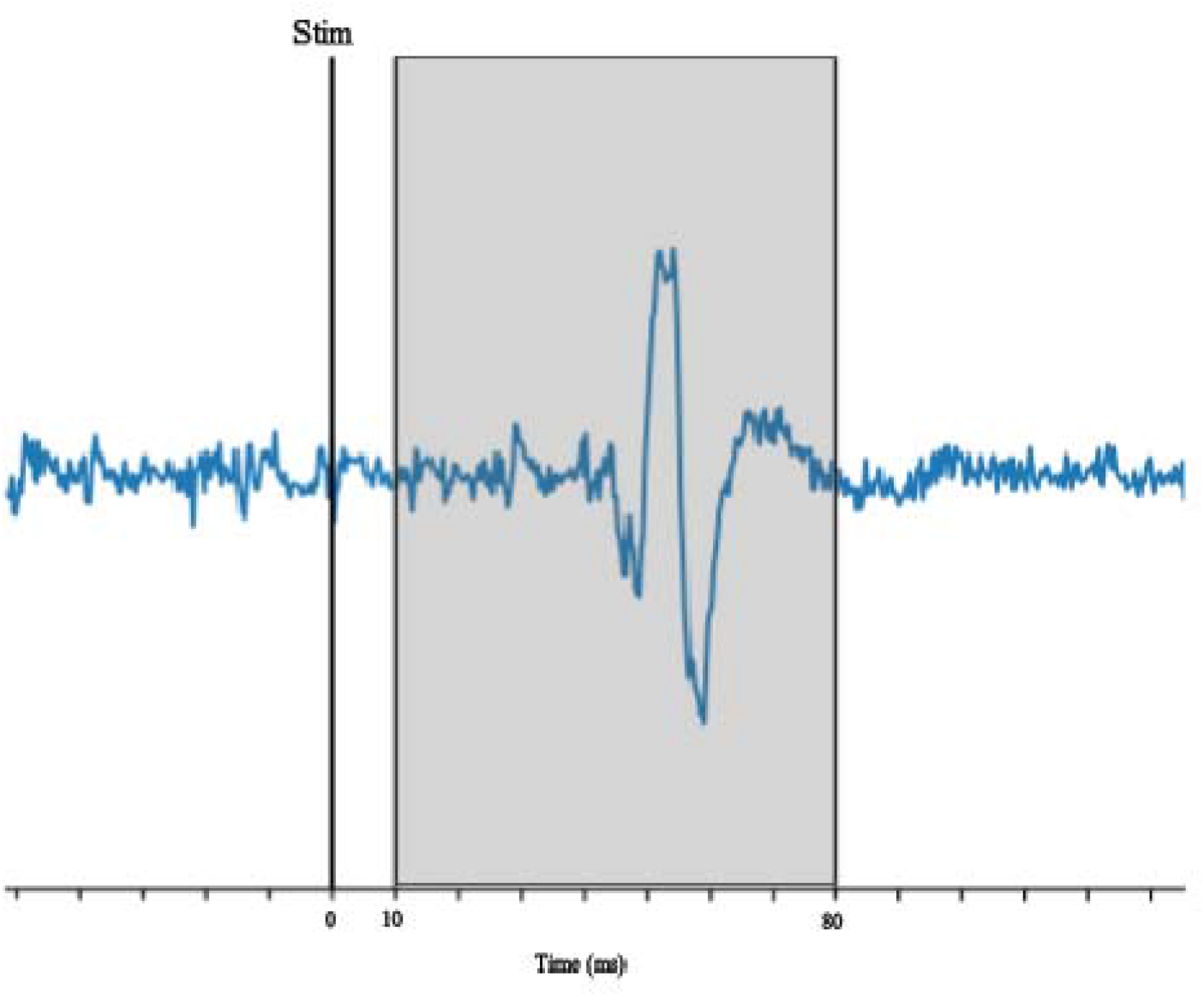
Example of extracted MEP. The blue line represents the recorded EMG signal. The detected stimulation is represented by the vertical black line. MEPs were extracted using a post-stimulus window of 10-80 ms following each detected stimulation, as illustrated by the shaded region.

## Results

### MEP Reliability in Target Muscle

The first analyses focused on evaluating the reliability of mean rectified magnitude and peak-to-peak amplitude of MEP size in the target muscle, i.e., the right TA, at rest. To test for session differences in each dependent measure, separate analyses for each quantification of MEP size compared a linear mixed-effects regression model fitted with individual differences treated as a random factor across participants and including the fixed factors of session (1, 2, or 3), stimulation intensity (110% RMT, 120% RMT), and coil accuracy (distance of coil from target, reported in mm) with a null model (i.e., without the fixed effect of session). For both quantification methods, a likelihood ratio test showed that the model including session as a predictor fit the data significantly better than the null model (mean rectified magnitude: *χ*2(2) = 19. 56, *p <* . 001; peak-to-peak amplitude: *χ*2(2) = 32. 38, *p <* . 001), indicating a significant effect of session on MEP size. Given this, intraclass correlation coefficients (ICC) and the Bland-Altman analysis were used to further quantify inter-session reliability and variability.

#### Effect of stimulation intensity and quantification method on target-muscle reliability

In order to assess whether stimulation intensity (110% *vs*. 120% RMT) and quantification method (peak-to-peak amplitude *vs*. mean rectified magnitude) influenced the consistency of MEP size between sessions, ICC values were calculated across session for both measures of MEP size using the ICC(C,1) formula outlined by McGraw and Wong (1996). Specifically, ICC values were calculated for MEP size between sessions for each of the two stimulation conditions (110% and 120% RMT) for two different measures (mean rectified magnitude and peak-to-peak amplitude). An ICC value of 1 represents perfect repeatability of a measurement, with values greater than 0.6 indicating reasonably high repeatability [29]. The ICC values for the right TA are displayed in Table 3. There were significant ICC values observed across sessions for both stimulation conditions and quantification methods, with the highest reliability observed in the mean rectified magnitude quantification of MEP size recorded at 120% RMT (*ICC* = 0. 696, *p* < . 001). This result is similar to between-session reliability of MEP size in the TA at rest observed by Su et al. (2022), who reported an ICC value of 0.730 for MEPs recorded at 130% RMT.

To compare the variability of the stimulation intensities and quantification methods used to calculate MEP size, the coefficient of variation (CV) was calculated for MEP size between sessions for each of the two stimulation conditions (110% and 120% RMT) for both measures (mean rectified magnitude and peak-to-peak amplitude) using a ratio of the standard deviation to the mean. Figure 3 displays CV distributions calculated for each dependent measure (mean rectified magnitude and peak-to-peak amplitude, at both 110% and 120% RMT) in the right TA, such that the distribution of CV values for each individual session across subjects are plotted. Visual inspection of these distributions indicates that the lowest variability was observed using the mean rectified magnitude quantification of MEPs recorded at 120% RMT, as this distribution was skewed towards a CV value of zero, indicating lower variability reported across sessions and participants. This result is consistent with the ICC analysis, in which the numerically greatest reliability in MEP size was observed in the mean rectified magnitude quantification of MEPs recorded at 120% RMT as well.

**Figure 3:**
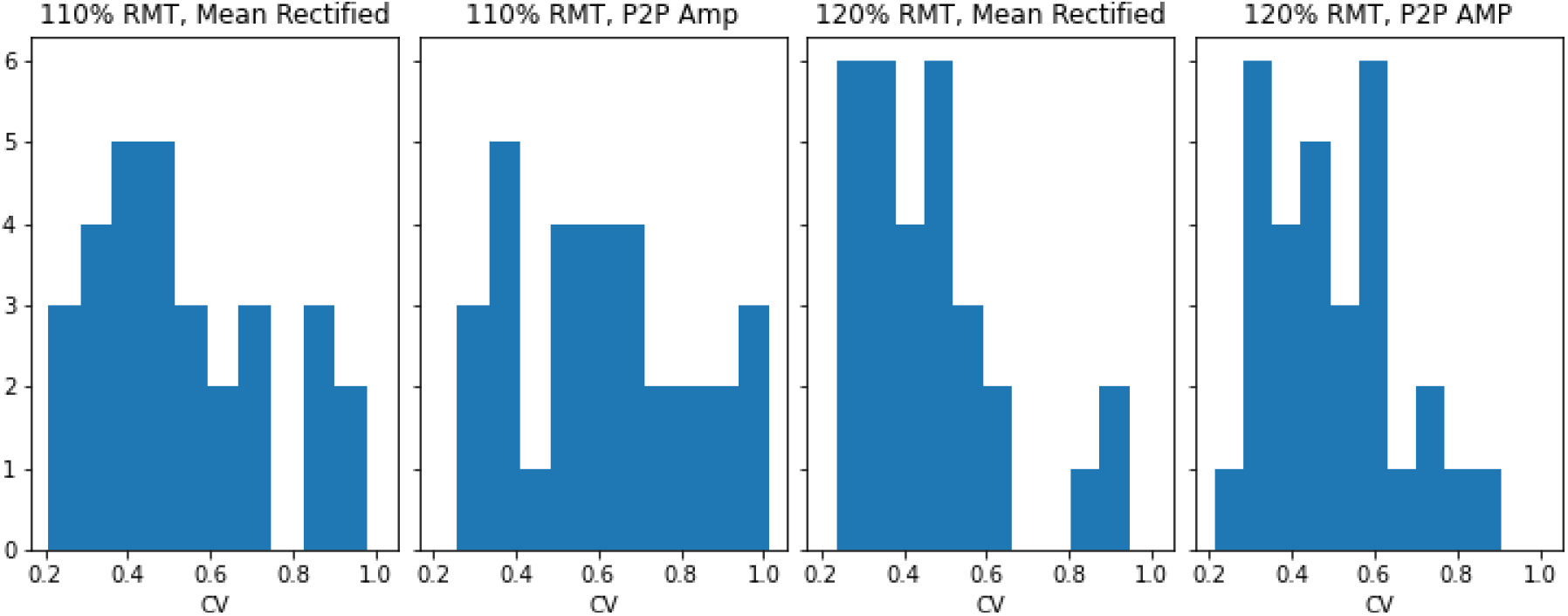
Coefficient of variation distributions across both dependent measures (mean rectified magnitude and peak-to-peak amplitude) recorded at both stimulation conditions (110% and 120% RMT) in the right TA. Bars represent cumulative sessions across participants with specific CV values. Distributions skewed towards CV values of zero indicate an overall lower amount of variability across all participants and sessions.

Due to observed differences in MEP size between sessions, Bland-Altman analysis was used to determine a baseline range of variability of MEP size in each muscle at rest. This analysis measures the degree of agreement between quantitative measurements by producing limits of agreement (i.e., a range of variability) using calculated means and standard deviations of the differences between two measurements [30]. In this case, the pairs of measurements were the mean MEP size of all combinations of sessions (i.e., sessions 1 and 2; sessions 2 and 3; sessions 1 and 3) for each dependent measure (peak-to-peak amplitude and mean rectified magnitude, both at 110% and 120% RMT). Figure 4 demonstrates the Bland-Altman analysis of MEP size in the right TA. The ranges of variability calculated for each dependent measure in the right TA are displayed in Table 1, which are used to quantify an expected range of variability between sessions without an effect of training.

**Table 1:**
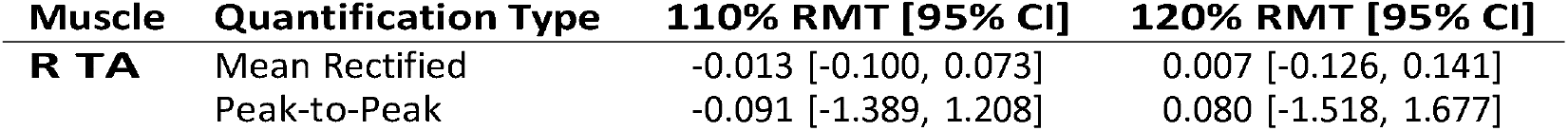
Bland Altman analysis in right TA. Overall mean difference in MEP size between sessions (mV).

**Figure 4:**
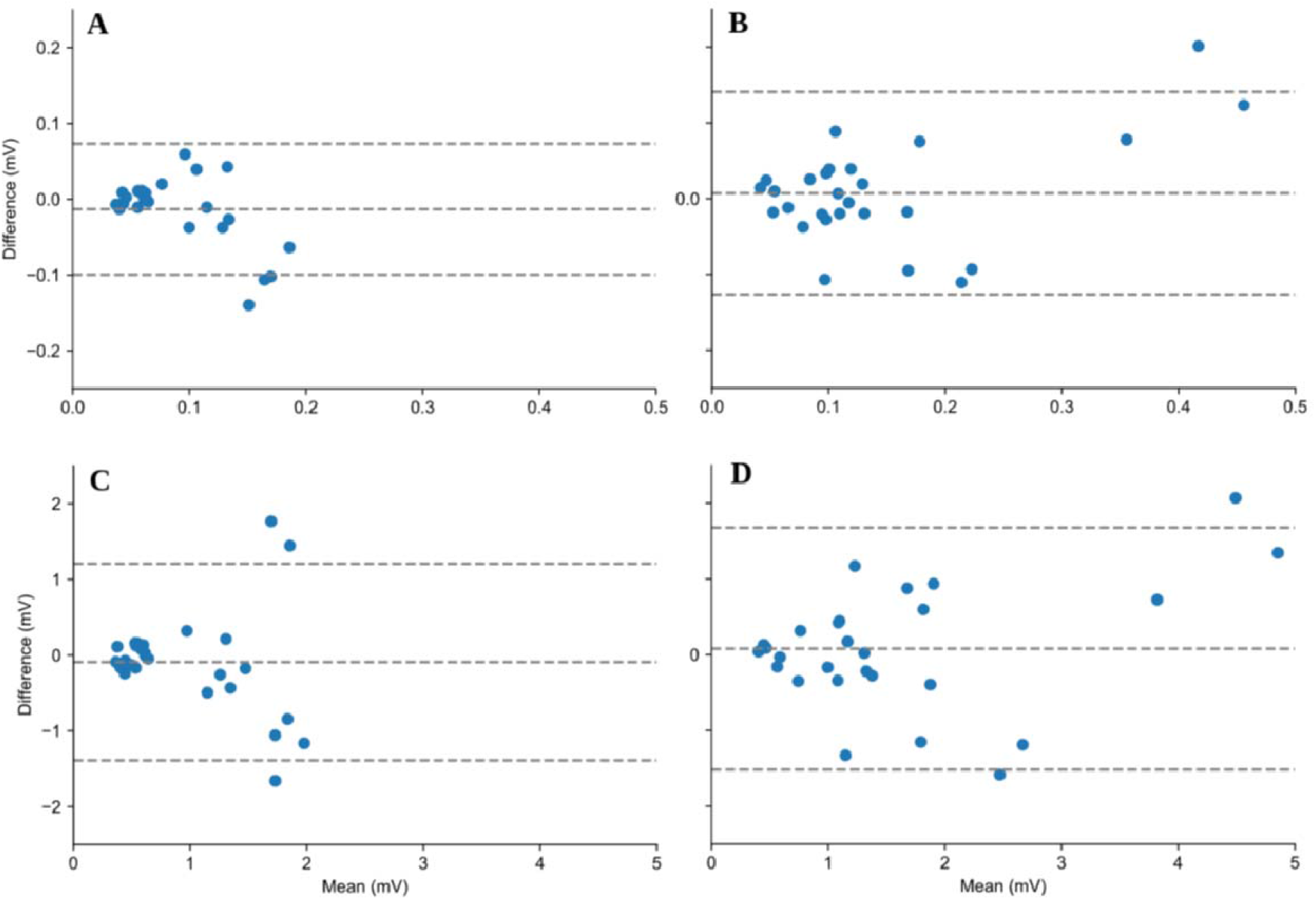
Bland-Altman analysis across subjects in the right TA. In each panel, the center dotted-line indicates the overall mean difference between measurements. The upper and lower bounds (i.e., dotted lines on the figure) represent 95% confidence intervals of the mean difference, which constitutes the range of variability expected for each dependent measure. Mean rectified magnitude quantification is displayed in panels A and B at 110% RMT and 120% RMT, respectively. Peak-to-peak amplitude quantification is displayed in panels C and D at 110% RMT and 120% RMT, respectively.

### MEP Reliability in Non-Target Muscles

Given the highest reliability in the target muscle (right TA) was observed at 120% RMT when quantified using the mean rectified magnitude, these parameters were used to assess MEP reliability in the remaining seven non-target muscles (left TA; right and left soleus, rectus femoris, biceps femoris). Following the analysis procedure outlined for the right TA, non-target muscles were assessed for session differences for mean rectified MEP magnitude. As stated above, separate analyses for each quantification of MEP size compared a linear mixed effects regression model fitted with individual differences treated as a random factor across participants and including the fixed factors of session (1, 2, or 3), stimulation intensity (110% RMT, 120% RMT), and coil accuracy (distance of coil from target, reported in millimeters) with a null model without the fixed effect of session. A significant effect of session on MEP size was observed in all non-target muscles: left TA (*χ*2(2) = 50. 28, *p <* . 001); right soleus (*χ*2(2) = 14. 98, *p <* . 001); left soleus (*χ*2(2) = 53. 91, *p <* . 001); right rectus femoris (χ (2) = 6. 09, *p* = 0. 047)); left rectus femoris (*χ*2(2) = 32. 50, *p <* . 001; right biceps femoris (*χ*2(2) = 21. 94, *p <* . 001; left biceps femoris (*χ*2(2) = 44. 87, *p <* . 001.

The ICC results for mean rectified MEP magnitude of all non-target muscles recorded at 120% RMT are displayed in Table 4. Poor reliability was observed in all non-target muscles according to the ranges outlined by McGraw & Wong (1996), with the exception of the right rectus femoris, in which moderate reliability was observed (ICC=0.518).Following the procedure outlined for the right TA, Bland-Altman analysis was also performed in order to quantify ranges of variability for all non-target muscles, as displayed in Table 2.

**Table 2:** Bland-Altman analysis of mean rectified MEP magnitude in non-target muscles. Overall mean difference in MEP size between sessions (mV).

| Muscle | 120% RMT [95% CI] |
| --- | --- |
| **R Soleus** | -0.001 [-0.083, 0.081] |
| **R Rectus Femoris** | 0.002 [-0.120, 0.204] |
| **R Biceps Femoris** | -0.0002 [-0.067, 0.066] |
| **L TA** | -0.009 [-0.158, 0.140] |
| **L Soleus** | -0.015 [-0.067, 0.037] |
| **L Rectus Femoris** | -0.005 [-0.105, 0.095] |
| **L Biceps Femoris** | 0.096 [-0.649, 0.842] |

**Table 3:**
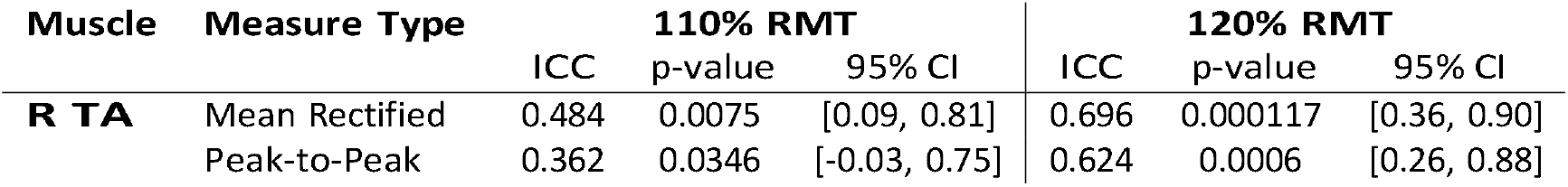
Intraclass Correlation Coefficient (ICC(C,1)) for EMG Measures at 110% and 120% RMT in the right TA.

| Muscle | Measure Type | 110% RMT |  |  | 120% RMT |  |  |
| --- | --- | --- | --- | --- | --- | --- | --- |
|  |  | ICC | p-value | 95% CI | ICC | p-value | 95% CI |
| **R TA** | Mean Rectified | 0.484 | 0.0075 | [0.09, 0.81] | 0.696 | 0.000117 | [0.36, 0.90] |
|  | Peak-to-Peak | 0.362 | 0.0346 | [-0.03, 0.75] | 0.624 | 0.0006 | [0.26, 0.88] |

**Table 4:** Intraclass Correlation Coefficient (ICC(C,1)) for mean rectified MEP magnitude of non-target muscles at 120% RMT.

| Muscle | ICC | p-value | 95% CI |
| --- | --- | --- | --- |
| **R Soleus** | 0.080 | 0.3220 | [-0.23, 0.55] |
| **R Rectus Femoris** | 0.518 | 0.0045 | [0.13, 0.83] |
| **R Biceps Femoris** | 0.142 | 0.2220 | [-0.19, 0.60] |
| **L TA** | 0.383 | 0.0275 | [-0.01, 0.76] |
| **L Soleus** | 0.204 | 0.1445 | [-0.15, 0.65] |
| **L Rectus Femoris** | 0.252 | 0.0986 | [-0.12, 0.68] |
| **L Biceps Femoris** | -0.040 | 0.5566 | [-0.30, 0.43] |

## Discussion

The current study assesses the reliability of MEPs elicited in multiple lower extremity muscles from a single TMS location over the cortex. Previous studies have reported high MEP reliability in lower extremity muscles when assessed individually, including the TA [8, 9, 10, 11], vastus medialis [12, 13], vastus lateralis [14, 15], and rectus femoris [16]. Although TMS has been shown to effectively elicit MEPs in multiple lower extremity muscles from a single location [20, 19], no studies have examined the reliability of MEPs generated bilaterally in muscles beyond the primary target muscle. To address this gap, we recorded MEPs from bilateral TA, soleus, rectus femoris, and biceps femoris muscles across three sessions separated by approximately one week. Stimulation was delivered at both 110% and 120% of each participant’s RMT, and MEP size was quantified using both mean rectified magnitude and peak-to-peak amplitude to compare reliability between these two methods.

The overall consistency of MEP size across conditions (mean rectified magnitude at 110% and 120% RMT, and peak-to-peak amplitude at 110% and 120% RMT) was assessed using ICC(C,1) as defined by McGraw and Wong (1996). In the right TA (i.e., target muscle for TMS), the highest consistency between sessions was observed at 120% RMT (ICC=0.696, p*<*0.001). This result is comparable to previously reported consistency of MEP size between sessions when targeting the TA at rest (ICC=0.73) [31]. When comparing the coefficients of variation between stimulation intensities and quantification methods in the right TA, mean rectified magnitude of MEPs recorded at 120% RMT reported the lowest variation, demonstrating that this condition was the most reliable in the target muscle. Other non-target muscles demonstrated lower consistency between sessions, and did not follow the pattern of reliability found in the target muscle. Additionally, the factor of session demonstrated a significant effect on MEP size for all recorded muscles, with the exception of the peak-to-peak amplitude quantification in the right rectus femoris.

Together, these patterns of results suggest that, despite TMS eliciting consistent MEP sizes across sessions in the target muscle at a higher stimulation intensity, there are demonstrated effects of session on MEP size across all muscles. Thus, the Bland-Altman analysis was used to quantify a range of variability of MEP size between sessions for each muscle and condition. These ranges will serve as a representation of expected variation in MEP size in unimpaired subjects without training intervention. Due to the depth of the representation of the lower limb in the motor cortex, a double-cone coil was used to enable the magnetic field generated by TMS to penetrate the cortex at the desired location [32]. Therefore, when targeting a single muscle of the lower limb, activity will be elicited in additional muscles due to the limited focality of the TMS magnetic field at this depth generating descending volleys of corticospinal pathways to these muscles [32]. The current study found co-activated muscles (e.g., soleus, rectus femoris, and biceps femoris) to demonstrate considerably lower between-session reliability in MEP size than the targeted muscle (TA). As described by Eisner-Janowicz et al. (2023), lower limb muscles demonstrate different levels of corticospinal excitability in unimpaired populations, which could be attributed to the differences in reliability observed here. Distal lower limb muscles (i.e., TA) exhibit higher excitability, and thus require less stimulation to generate an MEP than more proximal muscles of the lower limb [21]. In the current study, the RMT used to determine the stimulation level of TMS for each participant was contingent only on the response of the right TA. Thus, the stimulation intensity may not have been adequate to consistently produce MEPs of the same size in co-activated muscles, contributing to the higher variability observed in these muscles. Additionally, Eisner-Janowicz et al. (2023) reported that the biceps femoris muscle receives the weakest cortical input in comparison to the TA, soleus, and rectus femoris, which is consistent with the left biceps femoris demonstrating the lowest consistency across sessions in the current study.

Although we observed variation in MEP size between sessions, especially in muscles co-activated with the target muscle, measurements of corticospinal excitability can still reasonably be assessed across sessions once an expected range of variation has been established in these muscles. Using the Bland-Altman analysis, the current study provides a quantified range of variability of MEP size between sessions without training intervention. This range not only demonstrates the expected day-to-day variability in the target muscle, but can also account for differences in excitability of co-activated muscles that may influence MEP size across sessions. This process is especially important when assessing changes in corticospinal excitability associated with motor learning, as providing a baseline effect of session on the variability of MEP size can increase confidence that any observed changes in MEP size that fall outside of this range would be due to the effect of training, rather than simply variability of the measurement.

